# The genome sequence of the avian vampire fly (*Philornis downsi*), an invasive nest parasite of Darwin’s finches in Galápagos

**DOI:** 10.1101/2021.06.09.447800

**Authors:** Melia Romine, Sarah A. Knutie, Carly M. Crow, Grace J. Vaziri, Jaime Chaves, Jennifer A.H. Koop, Sangeet Lamichhaney

## Abstract

The invasive avian vampire fly (*Philornis downsi*) is considered one of the greatest threats to the unique and endemic avifauna of the Galápagos Islands, Ecuador. The fly parasitizes nearly every passerine species, including Darwin’s finches, in the Galápagos. The fly is thought to have been introduced from mainland Ecuador, although the full pathway of invasion is not yet known. The majority of research to date has focused on the effects of the fly on the fitness of avian hosts and explorations of mitigation methods. A lag in research related to the genetics of this invasion demonstrates, in part, a need to develop full-scale genomic resources with which to address further questions within this system. In this study, an adult *P. downsi* collected from San Cristobal Island within the Galápagos archipelago was sequenced to generate a high-quality genome assembly. We examined various features of the genome (e.g., coding regions, non-coding transposable elements) and carried out comparative genomics analysis against other dipteran genomes. We identified lists of gene families that are significantly expanding/contracting in *P. downsi* that are related to insecticide resistance, detoxification, and potential feeding ecology and counter defense against host immune responses. The *P. downsi* genome assembly provides an important foundational resource for studying the molecular basis of its successful invasion in the Galápagos and the dynamics of its population across multiple islands. The findings of significantly changing gene families associated with insecticide resistance and immune responses highlight the need for further investigations into the role of different gene families in aiding the fly’s successful invasion. Furthermore, this genomic resource will also better help inform future research studies and mitigation strategies aimed at minimizing the fly’s impact on the birds of the Galápagos.

## Introduction

The invasive avian vampire fly (*Philornis downsi*) (Figure 1a) is considered among the greatest threats to the unique and endemic avifauna of the Galápagos Islands, Ecuador (Causton *et al.*, 2013). The fly parasitizes eleven species of Darwin’s finches (Figure 1b) as well as nearly every other Galápagos passerine species (Causton *et al.*, 2013; Fessl *et al.*, 2018). The earliest record of adult flies from the Galápagos occurred in 1964, but the fly was not observed in the nests of birds until 1997 (Fessl and Tebbich, 2002; Causton *et al.*, 2006). The fly has now been found on all major islands within the archipelago except Española, Genovesa, Darwin, and Wolf (Causton *et al.*, 2013). The impact of parasitism by the fly has been severe in some populations of birds in Galápagos and while the effects are variable, some studies have reported near-total nest failure rates due to parasitism (Dudaniec *et al.*, 2007; Koop *et al.*, 2011; Koop, Le Bohec, *et al.*, 2013; O’Connor *et al.*, 2014; Knutie *et al.*, 2016a; Heimpel *et al.*, 2017; Addesso *et al.*, 2020). The fly has also been implicated in the decline of the medium tree finch (*Camarhyncus pauper*), the warbler finch (*Certhidia olivacea*), and the mangrove finch (*Camarhyncus heliobates*) (Dvorak *et al.*, 2004; Grant *et al.*, 2005; Cunninghame *et al.*, 2017; Peters and Kleindorfer, 2018; Bulgarella *et al.*, 2019). Furthermore, the potential for population-level declines of even relatively prominent bird species, e.g., the medium ground finch, have also been demonstrated using predictive models (Koop *et al.*, 2015).

**Figure 1:**
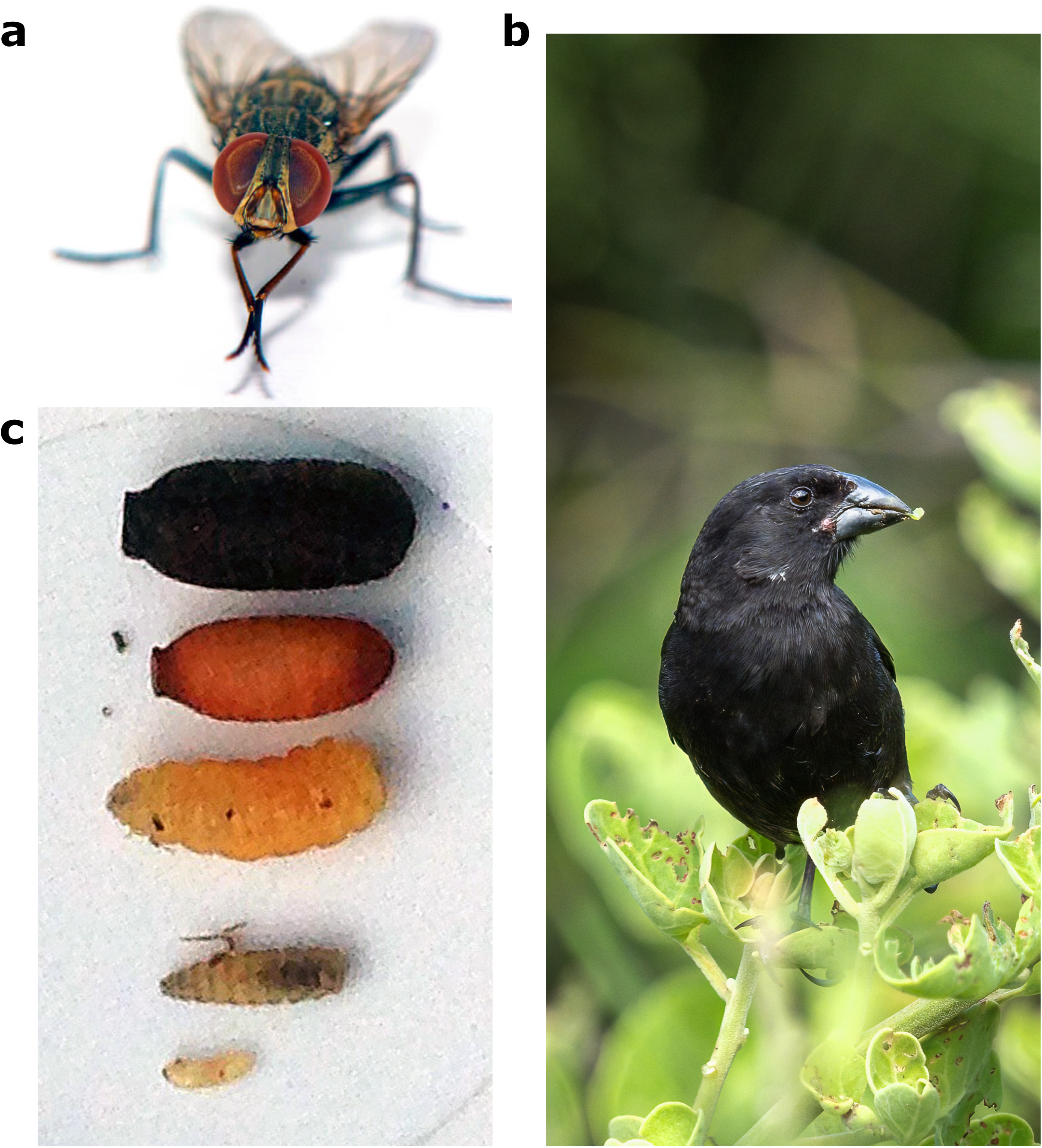
The avian vampire fly, Philornis downsi **(a)**, parasitizes many endemic bird species of the Galapagos Islands, including the medium ground finch, Geospiza fortis **(b)**. The fly is parasitic in its larval forms **(c**, bottom three) when it feeds on the blood and other fluids of its avian hosts. The larva then pupates **(c**, second from top) and eclose **(c**, top) as adult flies.

The genus *Philornis* includes approximately 50 species found primarily in the Neotropics and into North America (Dodge, 1955, 1963; Spalding *et al.*, 2002; Dudaniec and Kleindorfer, 2006; Couri *et al.*, 2007). Of the described species within *Philornis*, only two species (including *P. downsi*) are free-living ectoparasites within the nests of their hosts (Fessl *et al.*, 2006), while most others are subcutaneous on their hosts. Several *Philornis* species are found in mainland Ecuador (Bulgarella *et al.*, 2015, 2017), but *P. downsi* remains the only recorded species of the genus present, to date, in the Galápagos. *P. downsi* is thought to have been introduced from mainland Ecuador, though the full pathway of invasion is not yet known (Fessl *et al.*, 2018). Preliminary population genetics studies show that flies within the archipelago have a high degree of relatedness relative to those on the mainland, which supports the hypothesis of a relatively recent invasion and also the possibility of continued gene flow between populations in the Galápagos (Dudaniec *et al.*, 2008; Koop *et al.*, 2021).

A collaborative research effort has been made to continue monitoring the effects of the avian vampire fly on bird populations in the Galápagos. These efforts aim to identify source population(s), the pathway and mechanism of invasion, and possible long-term mitigation methods, including the Sterile Insect Technique and biocontrol (Causton *et al.*, 2013, 2019). Current stopgap mitigation efforts in the Galápagos include the direct application of insecticide to bird nests. Newly hatched nestlings are removed, and a permethrin solution is sprayed inside the nest and allowed to dry, at which point the nestlings are placed back in the nest. This method has been shown to effectively reduce parasites in the nest and increase fledging success (Koop *et al.*, 2011). A continuation of this method relies on birds incorporating permethrin-treated cotton into their nests, achieving similar increases in fledging success (Knutie *et al.*, 2014). Both methods, with subtle variations, are being used in attempts to increase the nesting success of the critically endangered mangrove finch, currently one of the most endangered birds in the world (Fessl *et al.*, 2018). The same method is also being used successfully to protect endangered bird species against their respective parasites in Australia (Alves *et al.*, 2020).

Despite the robust number of studies that explored the effects of these flies on hosts in the Galápagos, questions remain about the underlying ecological and evolutionary mechanisms of their successful invasion in the Galápagos. This knowledge gap demonstrates the need to develop a full-scale genomic resource of the fly as it provides a critical knowledge base from which to explore these questions, similar to other parasites of concern (Scott *et al.*, 2020). In this study, we generated a high-quality draft genome of the avian vampire fly, which is expected to become an important resource of future molecular studies in this system. We further carried out comparative genomics analysis with additional published dipteran genomes that identified evidence of significantly changing gene families associated with insecticide resistance, detoxification, and possible counter defense against host immune responses. These results serve as the first step toward investigations of this fly’s ability to rapidly evolve traits associated with its successful invasion in the Galápagos. From an applied aspect, these genomic resources will also help inform future research studies and mitigation strategies aimed at minimizing the fly’s impact on the birds of the Galápagos.

## Materials and Methods

### Sampling and DNA extraction for genome sequencing

Flies were collected in Jardín de las Opuntias on San Cristobal Island, Galápagos, Ecuador (−0.9491651°, −89.5528454°) in March-April of 2019. Adult flies were reared from pupae collected in the nests of small ground finches (*Geospiza fuliginosa*). When the nests were empty because nestlings died or fledged, all larvae and pupae were extracted from the nest and placed in a falcon tube with a modified lid that allowed airflow. After flies eclosed from their pupal case, they were placed in the freezer, then preserved in 95% ethanol. Preserved flies were transported to the University of Connecticut, then shipped to Northern Illinois University for further processing. DNA was extracted using DNeasy Blood and Tissue Kit (Qiagen, Valencia, California, USA) on whole fly samples after wings and legs were removed using forceps. All samples were treated with Monarch® RNase A (New England Biolabs) to remove RNA from genomic DNA samples. Multiple samples were extracted and those with low yields of DNA or contamination were discarded from further processing. Quantification of double-stranded DNA was done using QuBit® and samples were run on a gel to assess fragmentation.

### Estimate of genome size and ploidy

We prepared a 10X Chromium GEM library from the extracted DNA above according to the manufacturer’s recommended protocols. The library was sequenced using the Illumina 10X platform to generate paired-end 150 bp reads. We used a k-mer based approach to estimate genome size, heterozygosity, and repeat content from unprocessed short sequencing reads (Vurture *et al.*, 2017). We also used the k-mer distribution to extract heterozygous k-mer pairs to assess the ploidy level in *P. downsi* (Ranallo-Benavidez *et al.*, 2020). These estimates of genome size and ploidy were then used to choose parameters for the downstream genome assembly pipeline.

### Genome assembly and annotation

The 10x Chromium linked reads generated above were used to generate a reference genome using Supernova.v.2.1.1 (Weisenfeld *et al.*, 2017) with default parameters. The genome contiguity statistics such as scaffold N_50_ and the total number of scaffolds etc. were calculated using custom Perl scripts. We further compared the draft genome assembly against a set of conserved genes in Diptera using BUSCO (Waterhouse *et al.*, 2017) to assess gene-space completeness. We also mapped the short sequencing reads back to the draft genome assembly using BWA (Li and Durbin, 2009) to estimate how many short-sequence reads were used to build the reference genome and infer genome completeness. The annotation of the *P. downsi* genome was done using the MAKER annotation pipeline (Cantarel *et al.*, 2008). We combined protein homology evidence (using publicly available protein datasets from major dipteran lineages, downloaded from the Ensemble database (Cunningham *et al.*, 2019) and *ab-initio* gene predictions models to annotate the genome.

### Transposable elements (TE) in *P. downsi*

We combined homology-based and de novo approaches using Repeatmasker (Smit Hubley P., 2013; Flynn *et al.*, 2019) to characterize TEs in *P. downsi*. We used two different sets of repeat libraries, (a) reference repeat library downloaded from the Repbase database (Bao *et al.*, 2015) and (b) *de novo* custom-built species-specific repeat library generated for *P. downsi* using RepeatModeler (Flynn *et al.*, 2019). The usage of the *de novo* species-specific repeat library increased the accuracy of detection and annotation of transposable elements. For an unbiased comparison of repeat landscapes among all species, we used similar approaches to detect TEs in the Housefly *(M. domestica)*, Stable fly *(S. calcitrans)*, Tsetse fly *(G. morsitans)*, Mediterranean fruit fly *(C. capitata)*, Fruit fly *(D. melanogaster)* and Yellow fever mosquito *(A. aegypti)*, together with an outgroup, Postman butterfly *(H. melpomene)*.

### Orthologs to other dipteran genomes

We inferred orthogroups and orthologs by comparative analysis of proteomes in *P. downsi* and six additional dipteran species (*M. domestica, S. calcitrans, G. morsitans, C. capitata, D. melanogaster, and A. aegypti)*, together with an outgroup, *H melpomene.* This analysis used the draft proteome of *P. downsi* generated from our genome annotation pipeline described above. Complete proteomes of each of the remaining six species were downloaded from the Ensemble database (Cunningham *et al.*, 2019). The orthologs inference was carried out using Orthofinder (Emms and Kelly, 2015).

### Gene family evolution

To explore the evolution of gene families in *P. downsi* to other Dipterans, we first constructed a maximum likelihood phylogeny using 3,070 single-copy orthologs among these eight species used for inferring orthologues (Figure 3a). We then used this phylogeny to analyze the changes in gene family size across lineages leading to each of these eight species using a maximum likelihood approach (CAFÉ) (De Bie *et al.*, 2006) that uses a birth and death process to model gene gain and loss across a phylogenetic tree.

## Results

### Genome sequencing and assembly

We sampled an adult *P. downsi* fly in Jardín de las Opuntias on San Cristobal Island, Galápagos. DNA library was prepared using 10X Chromium linked-read approach (Zheng *et al.*, 2016) and sequenced using Illumina 10X platform to generate paired-end 150 bp reads which resulted in ~479 million read pairs (~72 Gb of raw sequence data). We first generated a k-mer distribution based on these short-sequencing reads for preliminary characterization of genome structure in *P. downsi* (e.g., genome size, an abundance of repetitive elements, rate of heterozygosity) (Supplementary Table 1, Supplementary Figure 1) that further allowed us to make informed decisions on parameters needed for building a reference genome. We also utilized the k-mer distribution to estimate the ploidy level in *P. downsi*, which indicated it to be a diploid species (Supplementary Figure 2).

Based on genome size estimates using k-mer distributions from short sequence reads (Supplementary Table 1), we had ~ 91X sequence coverage for generating the draft *de-novo* genome assembly. The total estimated length of the draft assembly was 971.6 Mb. The assembly contained 41,176 total scaffolds (minimum 1000 bp to maximum 8.6 Mb) with scaffold N_50_ of 1.3 Mb. We further assessed genome contiguity and gene-space completeness using benchmarking universal single-copy orthologs (BUSCO) (Waterhouse *et al.*, 2017). Among 3,258 genes highly conserved across Diptera, 3,147 (95.8%) full-length genes were detected in the *P. downsi* draft genome assembly. Partial sequences of 71 genes were identified (2.2%) and only 67 (2.0%) were missing, indicating a high degree of contiguity in the *P. downsi* genome. In addition, 99.16% of total short sequencing reads aligned back to the draft genome indicating a high degree of genome completeness as well.

### Genome annotation

We combined protein homology-based evidence and *ab-initio* gene prediction models using the MAKER genome annotation pipeline (Cantarel *et al.*, 2008) which identified 15,774 protein-coding genes in the *P. downsi* genome. We compared the genome annotation of *P. downsi* with two other published and highly curated dipteran genomes, *Musca domestica* (Scott *et al.*, 2014) and *D. melanogaster* (Adams *et al.*, 2000). We annotated fewer genes in the *P. downsi* genome (15,774) compared to that of *M. domestica* (17,283) or *D. melanogaster* (17,468) (Table 1). The average length of genes is similar among *P. downsi* and *D. melanogaster*, whereas *M. domestica* typically has longer genes. The mean number of exons per gene is fewer in *P. downsi* compared to the other two species.

**Table 1:** Statistics of genomic features among three fly genomes.

| *Genome statistics* | *P. downsi* | *M. domestica*<br>(Scott et al. 2014) | *D. melanogaster*<br>(Adams et al. 2000) |
| --- | --- | --- | --- |
| Genome size (Gb) | 0.97 | 0.69 | 0.18 |
| Total genes | 15,774 | 17,283 | 17,468 |
| Avg. gene length (bp) | 4,789 | 13,747 | 5,830 |
| Total exons | 56,595 | 123,831 | 188,405 |
| Avg. exon length (bp) | 367 | 453 | 485 |
| Number of Exons per gene | 3.59 | 7.16 | 10.79 |

### Transposable elements in *P. downsi*

Mobile transposable elements (TE) are key features of eukaryotic genomes, being major determinants of genome size variation (Kapusta *et al.*, 2017; Lamichhaney *et al.*, 2021), and important contributors to the evolutionary potential of an organism (Pourrajab and Hekmatimoghaddam, 2021). We characterized the transposable elements in the *P. downsi* genome using homology-based (Smit Hubley P., 2013) and *de-novo* approaches (Flynn *et al.*, 2019). More than half of the genome (51.7%) of *P. downsi* consists of transposable elements (Supplementary Table 2). Among these sequences, 9.3% of the genome are retroelements (7.7% LINEs and 1.6% LTRs) and 23.4% DNA transposons. Short interspersed nuclear elements (SINEs), a major category of retroelements, were not detected in the *P. downsi* genome.

We also used similar methods to detect, annotate, and compare the repeat content across several other species including the house fly *(Musca domestica)*, stable fly *(Stomoxys calcitrans)*, tsetse fly *(Glossina morsitans)*, Mediterranean fruit fly *(Ceratitis capitata)*, fruit fly *(Drosophila melanogaster)* and yellow fever mosquito *(Aedes aegypti)*. The postman butterfly *(Heliconius melpomene*) was used as an outgroup. Transposable elements are known to be highly correlated with genome size across the tree of life (Kidwell, 2002; Lynch, 2007), and our results across various dipteran genomes are consistent with this pattern (Figure 2a).

**Figure 2:**
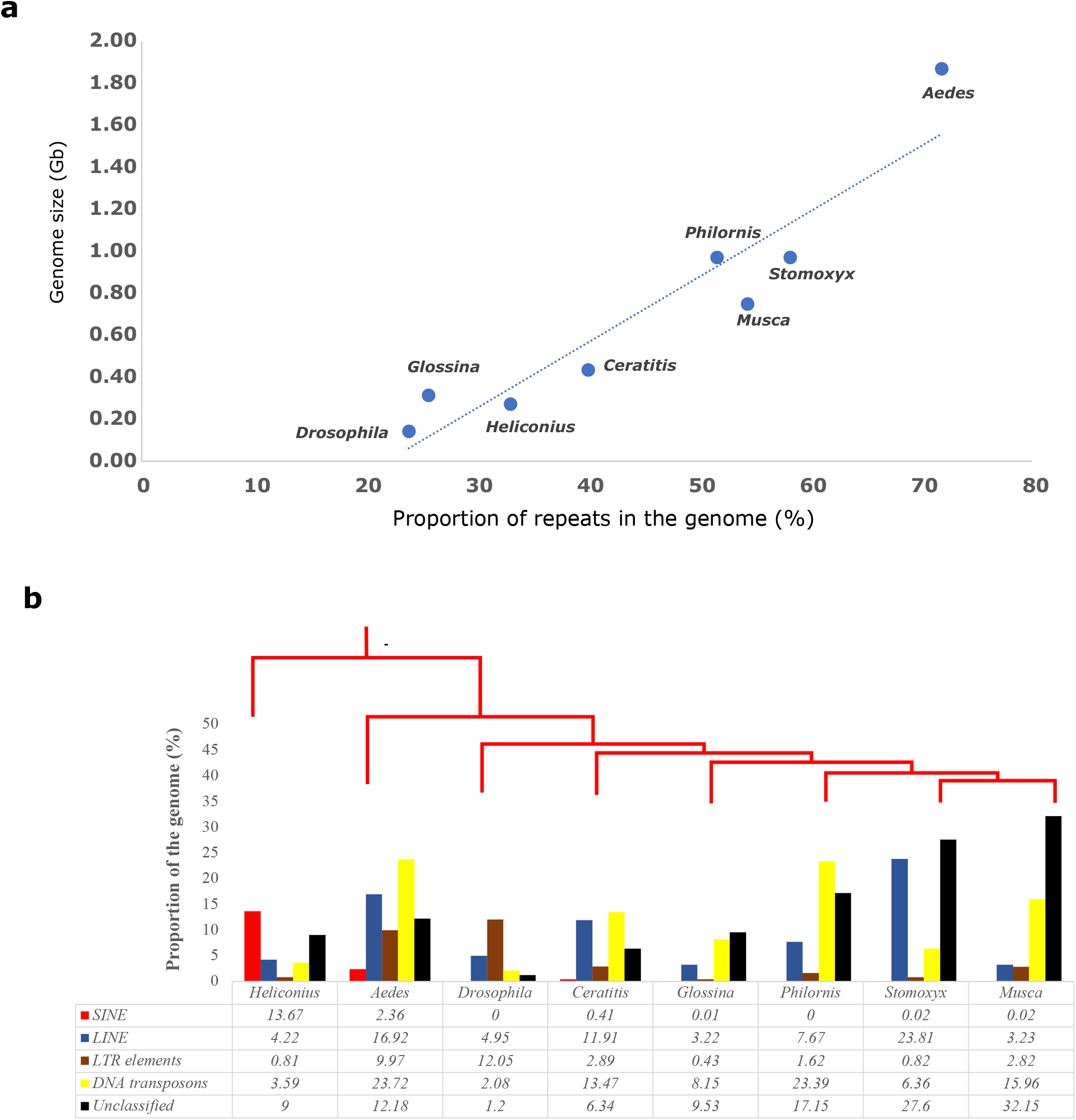
Landscape of Transposable elements in *P. downsi* **(a)** Comparison of repeat content and genome size across Diptera and its outgroup **(b)** Repeat statistics on various classes of transposable elements across dipteran genomes.

We also compared different categories of transposable elements across these seven genomes (Figure 2b). *P. downsi* had the highest proportion of DNA transposons (23.4%) among all taxa analyzed. In comparison, only 6.36% of the *S. calcitrans* genome and 15.96% of the *M. domestica* genome consisted of DNA transposons. Although *P. downsi* and *S. calcitrans* have similar genome sizes, *S. calcitrans* had slightly higher amounts of repeat content in its genome (58.3%), compared to *P. downsi* (51.7%) (Figure 2b). *S. calcitrans* had higher amounts of long interspersed nuclear elements (LINEs) (23.8%), compared to *P. downsi* (7.7%). We observed that most Diptera (except *Aedes*) have only a few SINE elements, one of the major classes of transposable elements (Figure 2b). LTR elements were relatively common in *D. melanogaster*, whereas other dipterans had a low amount of LTR elements (including *P. downsi*).

### Orthologs to other dipteran genomes and outgroup *Heliconius*

Comparative genomics analysis of 113,047 protein sequences from *P. downsi* and seven other species (listed above) identified 11,112 orthogroups. A total of 95,567 proteins (out of 113,047, 86.3%) were assigned to these orthogroups. The mean size of an orthogroup was 8.8 genes/species, and 3,069 orthogroups had singly copy genes in each species. A total of 5,754 orthogroups were shared among all eight species. Only a few orthogroups (minimum 496 in *A. aegypti* and a maximum of 1,445 in *M. domestica*) were present in fewer than four species (Figure 3a). The number of unique orthologs in each species was consistent with their phylogenetic relationships (e.g., *A. aegypti* and the outgroup *H. melpomene* had the highest number of unique orthogroups). The number of shared orthologs among each pair of species is presented in Supplementary Table 3. The number of orthologs identified in *P. downsi* is consistent with other published dipteran genomes (Figure 3a, Supplementary Table 3), indicating no major bias in the genome assembly and annotation pipeline used in this study.

**Figure 3:**
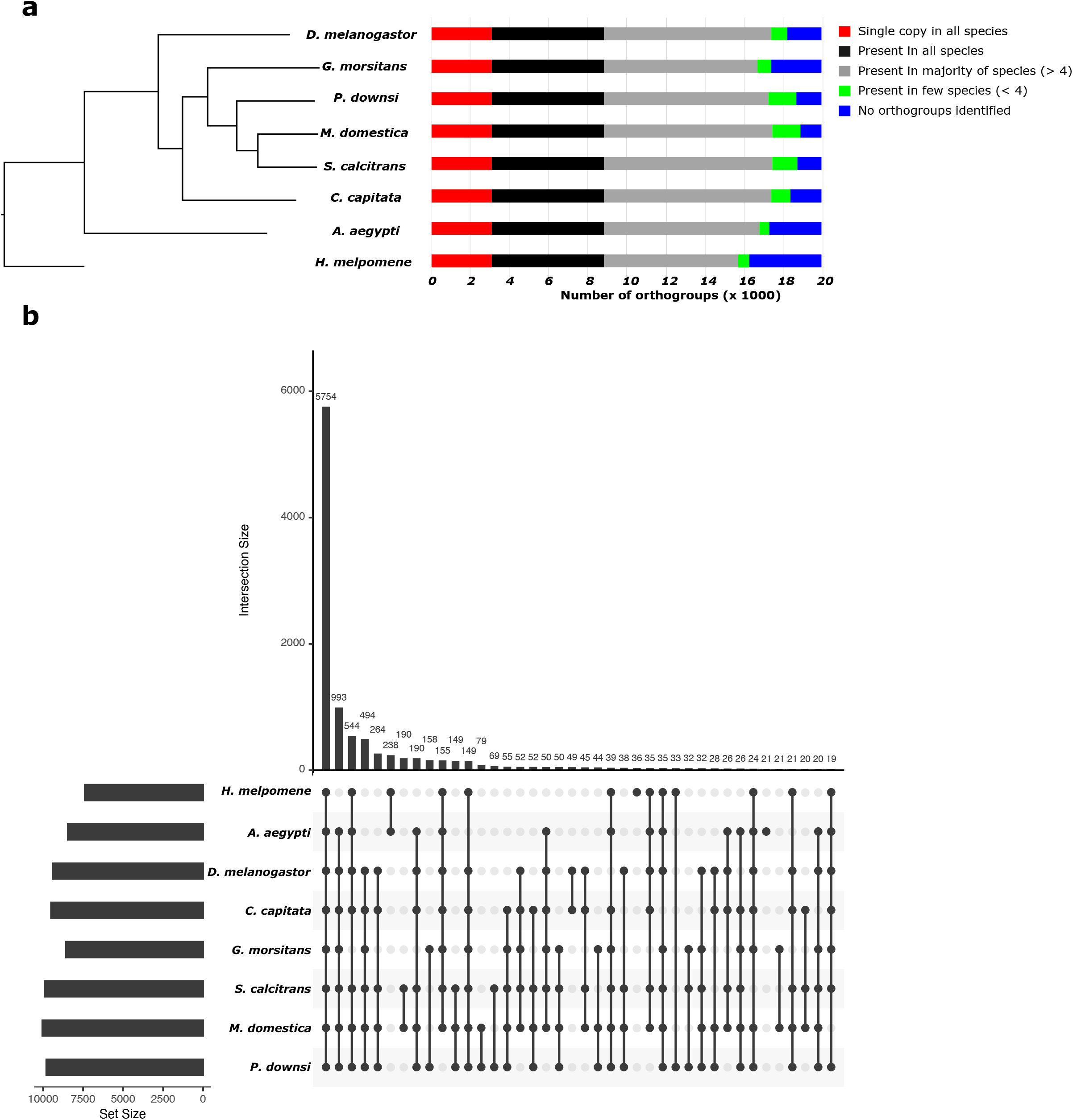
Orthogroups in *P. downsi* **(a)** Phylogenetic relationship between *P. downsi* and other seven published Diptera genomes, estimated using alignments from 3,069 orthogroups had singly copy orthogroups in each species. Horizontal bars for each species show number of orthogroups that are single-copy orthologs in all species, present in all species, present in the majority of species, present in few species, and unique to the species **(b)** Number of shared orthologs among all species.

Within *P. downsi*, 13,706 out of 15,774 (86.9%) annotated genes were assigned to orthogroups. We expect the remaining missing genes to either be the most recently evolved orphan genes in the branch leading to the *P. downsi* lineage or the consequence of a lack of inclusion of enough closely related species of *P. downsi* in the analysis. The distribution of these unique genes in the *P. downsi* genome is random and does not show specific clustering patterns across various locations of the genome.

We also compared the number of pairwise orthogroups that are uniquely shared among all eight species (Figure 3b). A total of 993 orthogroups were unique to Diptera (after excluding the outlier *H. melpomene)*, 79 orthogroups were shared only between *M. domestica* and *P. downsi* and 12 putative gene families were unique to only *P. downsi.* These 12 gene families consisted of 27 genes and the great majority had the best BLAST hits against “uncharacterized” or “hypothetical” proteins in other related species (Supplementary Table 4). This result indicates that these gene families that appeared “unique” in *P. downsi* are likely due to the lack of proper gene annotation in other species.

We further examined the gene ontology terms of the 993 orthogroups unique to Diptera using the PANTHER gene ontology database (Mi *et al.*, 2021). The common biological processes of these genes included localization, locomotion, immune system processes, response to stimulus, and reproduction (Supplementary Figure 3), many of which are likely key genes for the overall development and function of dipterans.

### Gene family evolution

We used a maximum likelihood approach to analyze changes in gene family size among the same eight species used for inferring orthologs (De Bie *et al.*, 2006). This method uses a statistical approach to model gene gain and loss, accounting for phylogenetic history and assess the significance of the observed gene family size differences among taxa. Across 11,112 orthogroups we identified before among eight taxa; 101 gene families were significantly expanding/contracting (p < 0.01) across the phylogenetic tree. 25 out of these 101 gene families were identified in the branch leading to *P. downsi*. The list of these gene families in *P. downsi* included those associated with insecticide resistance or detoxification and host defense or immunity proteins (Table 2).

**Table 2:** *List of* significantly expanding/contracting (p < 0.01) *gene families in P. downsi*

| Functional Categories | Gene Family |
| --- | --- |
| Insecticide Resistance and Detoxification | Cytochrome P450 |
|  | Glutathione S Transferase |
|  | Cuticular Protein |
| Defense against host Immunity | Fibrinogen C-terminal Domain-Containing Protein |
|  | Scp Domain-Containing Protein |

### Gene families associated with insecticide resistance and detoxification

#### Cytochrome P450 gene family

We examined the number of genes in the cytochrome P450 (CYP450) family across the seven Diptera and the *H. melpomene* outgroup. *P. downsi and M. domestica* have an expanded CYP450 gene family in comparison to their most recent common ancestor with *G. morsitans* (family Glossinidae) (Figure 4a). For example, The *P. downsi* CYP450 family is composed of 102 genes in comparison to 66 in *G. morsitans.* An even greater level of expansion was observed in two other members of Muscidae *(M. domestica:* 143 genes and *S. calcitrans:* 193 genes). Compared to *G. morsitans*, the expansion of CYP450 genes is mainly found in CYP4, CYP6, and CYP28 genes (Figure 4b). The CYP6 subfamily in *P. downsi* is composed of 25 genes, almost doubling the number from the 14 genes present in *G. morsitans.* We also found an expansion in the CYP4 subfamily from 15 genes in *G. morsitans* to 31 genes in *P. downsi.* A similar expansion was seen in the CYP28 subfamily (4 genes in *G. morsitans* to 8 genes in *P. downsi*).

**Figure 4:**
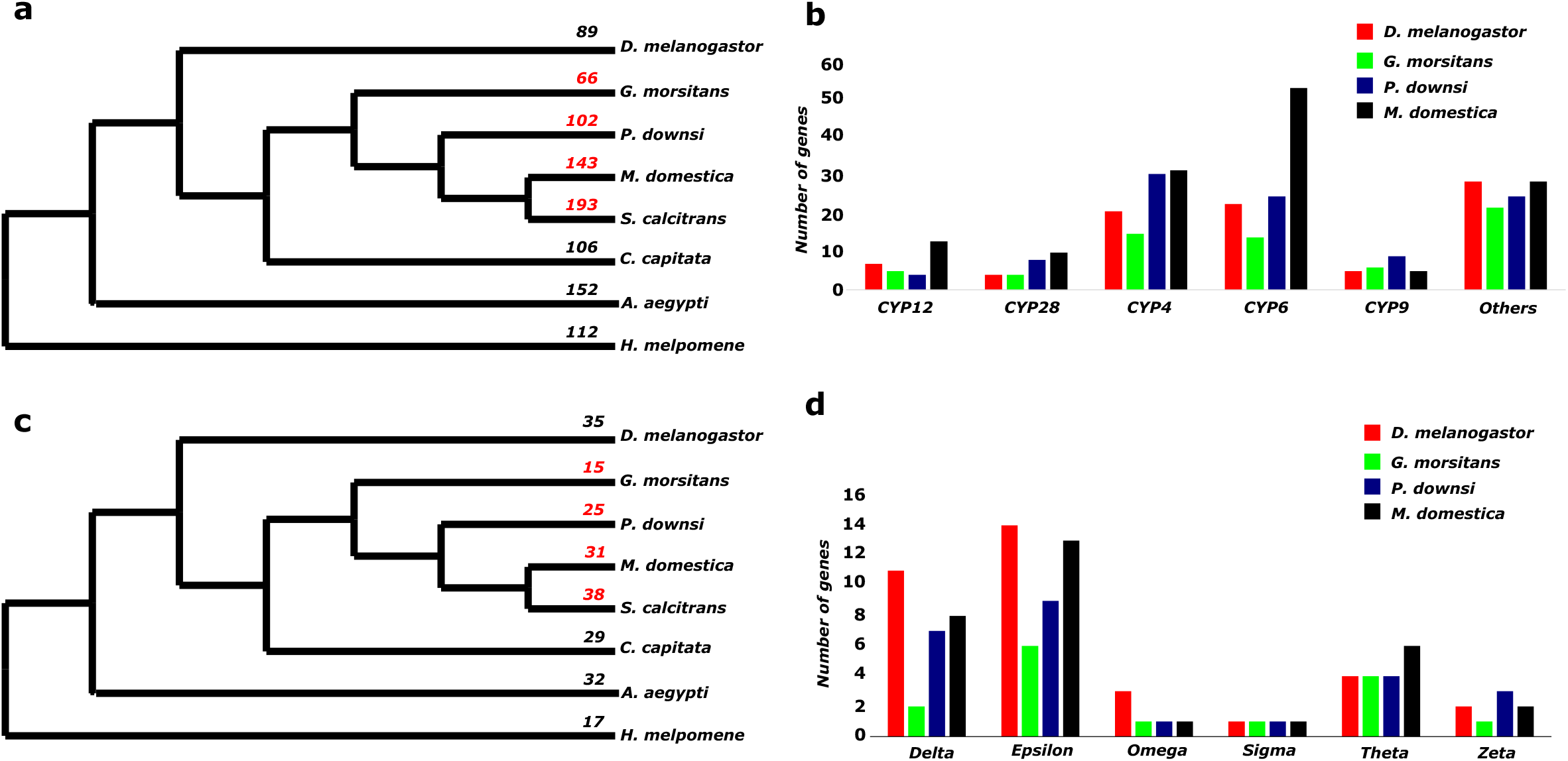
Gene family evolution **(a)** Number of genes in Cytochrome P450 gene family across Diptera **(b)** Number of genes in various subfamilies of Cytochromes P450 gene family in *G. morsitans* and *P. downsi, M. domestica* and *D. melanogaster* **(c)** Number of genes in Glutathione S-transferase gene family across Diptera **(d)** Number of genes in various subfamilies of Glutathione S-transferase gene family in *G. morsitans* and *P. downsi, M. domestica* and *D. melanogaster.*

#### Glutathione S-transferase gene family

The *P. downsi* Glutathione S-transferase family consists of 25 genes and the pattern across the dipterans is similar to observations of the Cytochrome P450 gene family expansion (Figure 4c). For example, in comparison to *G. morsitans* (15 genes), *P. downsi and M. domestica* have an expanded number of genes (25 genes in *P. downsi*, 31 genes in *M. domestica*). The GSTs are grouped into six subclasses (Delta, Epsilon, Omega, Sigma, Theta, and Zeta). The expansion of GSTs in *P. downsi* occurred mainly in the Delta, Epsilon, and Zeta subclasses (Figure 4d).

#### Cuticular Protein

The *P. downsi* cuticular gene family consists of 214 genes and the pattern across the dipterans is similar to observations of the Cytochrome P450 and GST gene family expansions (Supplementary Figure 4). For example, the *Muscidae* family (represented here by *P. downsi, M. domestica, and S. calcitrans)* has an expanded gene family in comparison to their most recent common ancestor with *G. morsitans* (115 genes). An even greater level of expansion was observed by two other members of *Muscidae (M. domestica* – 357 genes and *S. calcitrans* – 270 genes).

### Gene families associated with immunity

Two gene families (Fibrinogen C-terminal Domain-Containing Protein and Scp Domain-Containing Protein) associated with immunity were also identified as significantly changing in P. downsi (Table 2). Opposite of genes associated with insecticide resistance and detoxification (Cytochrome P450 and Glutathione S-transferase), these gene families showed a reduction in the number of genes in *P. downsi* compared to the most recent common ancestor with *G. morsitans* (Supplementary Table 5). They also show contraction in comparison with two included members of the Family Muscidae *(M. domestica and S. calcitrans).*

## Discussion

In this study, we report the first genome sequence of the avian vampire fly, a highly invasive parasitic nest fly that threatens endemic avifauna of the Galápagos Islands (Fessl *et al.*, 2018; Causton *et al.*, 2019). This genome is meant to serve as an important resource to research efforts aimed at characterizing the molecular mechanisms of the fly’s successful invasion in the Galápagos. The genome size of *P. downsi* is 971.6 Mb and its high quality is reflected both in terms of genome contiguity (scaffold N_50_ of 1.3 Mb) and completeness (98% BUSCO gene-space score). Interestingly, the total number of annotated genes in *P. downsi* is slightly lower compared to other published fly genomes (*M. domestica* and *D. melanogaster*) (Table 1). The low gene count likely reflects the lack of *P. downsi* specific transcriptome data in this study, which perhaps led to reduced gene predictions. We aim to improve the *P. downsi* genome annotations in the future using additional transcriptomic resources using additional flies collected from the Galápagos.

Transposable elements (TE) are typically non-coding sequences that can insert themselves in various places of the genome, often with neutral or deleterious phenotypic consequences (Bourque *et al.*, 2018). The role of TEs, as well as their evolution across insect genomes, is still an area of major research, but they are thought to be important drivers of genomic architecture depending on the location of the genome to which they insert themselves (i.e., coding versus non-coding regions). Furthermore, TE may also be a critical mechanism of adaptive evolution, as has been shown in an invasive ant species (Schrader *et al.*, 2014). Analysis of transposable elements in *P. downsi* and across other dipteran genomes showed a strong positive correlation between genome size and repeat content (Figure 2a), consistent with similar findings across other taxa (Lynch, 2007; Lamichhaney *et al.*, 2021). Interestingly, *P. downsi* had a higher number of DNA transposons (Class II TEs) than any other compared genome, including *M. domestica, S. calcitrans*, and *G. morsitans*. While long terminal repeat (LTR) transposons, LINEs, and SINEs) were present in the species studied, *P. downsi* had no SINEs, a finding consistent with a study by Petersen and colleagues (Petersen *et al.*, 2019) showing that SINEs contribute less than 1% to the TE content of dipterans. However, it is important to note that some SINEs may be present in *P. downsi* but are currently masked as unclassified. Future research should explore the role of TEs, especially DNA transposons, in aiding the invasion of *P. downsi* to the Galápagos.

Comparative genomics analysis of *P. downsi* and other additional dipteran genomes allowed us to identify gene families that were significantly expanded/contracted (p < 0.01) in *P. downsi*. The list of these gene families in *P. downsi* included those associated with insecticide resistance or detoxification and host defense or immunity proteins, and we predict that these gene families are associated with a successful invasion of *P. downsi* in the Galápagos (Table 1). E.g. CYP450 mono-oxygenases are a diverse superfamily of proteins, including enzymes, associated with catabolism and anabolism of xenobiotics and endogenous compounds. These monooxygenase-mediated metabolisms have allowed numerous insect species to develop insecticide resistance and detoxification (Scott, 1999; Wen *et al.*, 2011). The expansion of CYP450 genes in *P. downsi* is mainly found in CYP4 and CYP6 gene subfamilies (Figure 4b), which has also been shown in other species from the family Muscidae (Scott *et al.*, 2014; Olafson *et al.*, 2021). Interestingly, in higher Diptera, many of the genes within the CYP6 subfamily (e.g., CYP6A, CYP6G, CYP6D) are associated with insecticide resistance (Feyereisen, 2012). For example, the Cyp6g1 gene is involved in resistance to the insecticide Dichlorodiphenyltrichloroethane (DDT) in *D. melanogaster* (Festucci-Buselli *et al.*, 2005). The CYP4 subfamily can also influence the breakdown of synthetic insecticides (Iga and Kataoka, 2012). Cyp4d4v2, Cyp4g2, and Cyp6a38 can be co-up-regulated in-house flies that are resistant to the insecticide permethrin, which is used to control *P. downsi* in the Galapagos (Zhu *et al.*, 2008). Overall, the expansion of CYP4 and CYP6 subfamilies may indicate the evolution of insecticide resistance in *P. downsi* over macroevolutionary time (i.e., before arriving in the Galápagos), which could have facilitated its invasion to the Galápagos and might affect its management on the islands.

Glutathione S-transferase (GSTs) was another major expanding gene family in *P. downsi.* GSTs are a highly conserved, large family of dimeric enzymes associated with detoxification of endogenous and/or xenobiotic compounds, such as insecticides (Ketterman *et al.*, 2011). The GST family is further grouped into six subclasses (Delta, Epsilon, Omega, Sigma, Theta, and Zeta), with Delta and Epsilon being specific subclasses found in the class Insecta. We observed an expansion of GST genes in *P. downsi* relative to *G. morsitans*, but fewer than were found in *M. domestica and S. calcitrans*. The major expansions of the GST family in *P. downsi* were observed in Delta and Epsilon subclasses (Figure 4d). Insecticides, such as permethrin, are used to experimentally manipulate *P. downsi* abundance in bird nests to study the effects of the parasite on the health of the birds (Fessl *et al.*, 2010; Koop *et al.*, 2011; Koop, Le Bohec, *et al.*, 2013; Koop, Owen, *et al.*, 2013; Knutie *et al.*, 2014, 2016b; O’Connor *et al.*, 2014; McNew *et al.*, 2020; Addesso *et al.*, 2020). Increased expression of GSTs following permethrin exposure has been documented in several insect species including oriental fruit flies (*Bactrocera dorsalis*) (Hu *et al.*, 2008). Expansion of the GST gene family in *P. downsi* may be a result of such permethrin exposure. However, gene expression studies that further explore the role of the GST gene family in insecticide resistance and detoxification are needed across insect taxa including *P. downsi*. It is important to note that the expansion of CYP450 and GST families observed in *P. downsi* may be an artifact of phylogenetic relationships rather than ecological adaptations. Still, given the observed and predicted impacts of *P. downsi* on native endemic host populations, it is important to consider the implications of expanded gene families related to detoxification and insecticide resistance.

Previous studies have shown that Darwin’s finch species can increase specific antibody responses to parasitism (Koop, Owen, *et al.*, 2013). This response is most prominent in adult females that are likely parasitized while brooding nestlings or eggs on the nest. However, little is known about the ability of host bird immunological responses to effectively reduce fly fitness, in part, because so little is known about the fly itself. One of the most prominent questions is whether the *P. downsi* possesses the ability to counter defend against host immune responses. We identified a reduction in the size of two additional gene families (Fibrinogen C-terminal Domain-Containing and SCP domain-containing gene family), both with immune function properties in *P.downsi* (Table 1, Supplementary Table 5). Fibrinogen plays a key role in blood clot formation through the conversion of fibrinogen to insoluble fibrin (Weisel and Litvinov, 2017) and the C-terminal domain of fibrinogen is the primary binding site of platelets (Hanington and Zhang, 2011). The Sperm-coating glycoprotein (Scp) family contains, among other proteins, antigen 5 (Ag5), which is associated with the venom secretory ducts of stinging insects (Gibbs *et al.*, 2008). However, the interpretation of these results is difficult without further investigation into whether these genes are associated with innate immune responses of *P. downsi* toward their hosts or whether they might be important components of their feeding ecology. The goal of our study was not to make such inferences, but rather to highlight promising avenues of future research.

The invasion of *P. downsi* has had dramatic negative effects on the endemic avifauna of the Galápagos, including Darwin’s finches. As researchers work to better understand the pathway of invasion and the ecological and evolutionary processes that may have facilitated its invasion to the Galápagos, the need for a high-quality whole genome sequence has grown. The addition of this resource is therefore meant to provide the foundation for further investigations using genomics tools in this system. These genomic resources will further allow us to understand the evolution of the *P. downsi* defense in response to host defenses or disease resistance itself. Gene expression studies could shed light on the development of larvae and at what stage they are most vulnerable to host defenses. Further population-scale resequencing of various populations of *P. downsi* will also allow us to explore mechanisms of local adaptation of the parasite to the environment across islands.

## Data availability

All raw data generated in this study (raw short sequence reads and draft genome assembly) has been deposited to NCBI, under accession number PRJNAXXX. The final genome assembly and annotation can be found under the accession number GCA_XXX. Supplementary material is available on figshare.

## Acknowledgment

We would like to thank Lauren Albert, Taylor Verrett, and Corrine Arthur for field assistance and the Galápagos Science Center and the Galápagos National Park for logistical support. We also thank Noah K. Whiteman for his helpful comments on the manuscript. The sampling was done under Galápagos National Parks permits PC 28-19 and Genetic Access permit MAE-DNB-CM-2016-0041. This project was supported by the Department of Biological Sciences, Kent State University to SL, and the University of Connecticut to SAK.

## Author contributions

SL and JAHK conceived the idea. SAK, GJV, and JC collected the specimens. CMC and JAHK carried out the laboratory work. MR and SL designed and performed the bioinformatic analyses with the support of CMC and JAHK. SL and JAHK prepared the manuscript, and all authors edited and approved the final version.

## Supplementary Materials

**Supplementary Table 1:** Genome Statistics of P. downsi estimated from short-sequencing reads

| *Property* | *min* | *max* |
| --- | --- | --- |
| Heterozygosity | 1.40879% | 1.42194% |
| Genome Haploid Length | 796,797,589 bp | 799,050,413 bp |
| Genome Repeat Length | 232,405,721 bp | 233,062,812 bp |
| Genome Unique Length | 564,391,869 bp | 565,987,600 bp |
| Model Fit | 95.7713% | 98.7012% |
| Read Error Rate | 1.228% | 1.228% |

**Supplementary Table 2:** Repeat statistics in P. downsi genome

sequences: 41,176 total length: 971,6346,46 bp GC level: 35.10 % bases masked: 502,200,627 bp (51.69 %)
|  | number of<br>elements* | length<br>occupied | percentage<br>of sequence |
| --- | --- | --- | --- |
| ===== |  |  |  |
| Retroelements | 224657 | 90253146 bp | 9.29 % |
| **SINEs:** | **58** | **6761 bp** | **0.00 %** |
| Penelope | 3187 | 661022 bp | 0.07 % |
| **LINEs:** | **196268** | **74550945 bp** | **7.67 %** |
| CRE/SLACS | 0 | 0 bp | 0.00 % |
| L2/CR1/Rex | 43733 | 18550153 bp | 1.91 % |
| R1/LOA/Jockey | 15852 | 7502835 bp | 0.77 % |
| R2/R4/NeSL | 834 | 673762 bp | 0.07 % |
| RTE/Bov-B | 102395 | 29493616 bp | 3.04 % |
| L1/CIN4 | 1415 | 94740 bp | 0.01 % |
| **LTR elements:** | **28331** | **15695440 bp** | **1.62 %** |
| BEL/Pao | 12565 | 7084057 bp | 0.73 % |
| Ty1/Copia | 1561 | 775084 bp | 0.08 % |
| Gypsy/DIRS1 | 9951 | 7481940 bp | 0.77 % |
| Retroviral | 2481 | 143387 bp | 0.01 % |
| **DNA transposons** | **621139** | **227262774 bp** | **23.39 %** |
| hobo-Activator | 8749 | 1542960 bp | 0.16 % |
| Tc1-IS630-Pogo | 576750 | 212960682 bp | 21.92 % |
| En-Spm | 0 | 0 bp | 0.00 % |
| MuDR-IS905 | 0 | 0 bp | 0.00 % |
| PiggyBac | 1280 | 395680 bp | 0.04 % |
| Tourist/Harbinger | 4060 | 1924044 bp | 0.20 % |
| Other | 275 | 11451 bp | 0.00 % |
| Rolling-circles | 19653 | 4597029 bp | 0.47 % |
| **Unclassified:** | **828801** | **166508339 bp** | **17.14 %** |
| Total interspersed repeats: |  | 484024259 bp | 49.82 % |
| Small RNA: | 7148 | 2907047 bp | 0.30 % |
| Satellites: | 796 | 124775 bp | 0.01 % |
| Simple repeats: | 213281 | 8909635 bp | 0.92 % |
| Low complexity: | 34278 | 1638267 bp | 0.17 % |

**Supplementary Table 3:** Number of shared orthogroups among various dipteran species and their outgroup

|  | *H.melpomene* | *A.aegypti* | *C.capitata* | *S.calcitrans* | *M.domestica* | *P.downsi* | *G.morsitans* | *D.melanogaster* |
| --- | --- | --- | --- | --- | --- | --- | --- | --- |
| *H.melpomene* |  | 6,978 | 6,874 | 6,883 | 6,910 | 6,800 | 6,337 | 6,859 |
| *A.aegypti* | 6,978 |  | 8,015 | 8,008 | 8,018 | 7,796 | 7,228 | 7,995 |
| *C.capitata* | 6,874 | 8,015 |  | 9,123 | 9,186 | 8,855 | 8,014 | 9,096 |
| *S.calcitrans* | 6,883 | 8,008 | 9,123 |  | 9,650 | 9,197 | 8,087 | 9,041 |
| *M.domestica* | 6,910 | 8,018 | 9,186 | 9,650 |  | 9,295 | 8,144 | 9,117 |
| *P.downsi* | 6,800 | 7,796 | 8,855 | 9,197 | 9,295 |  | 8,103 | 8,770 |
| *G.morsitans* | 6,337 | 7,228 | 8,014 | 8,087 | 8,144 | 8,103 |  | 7,940 |
| *D.melanogaster* | 6,859 | 7,995 | 9,096 | 9,041 | 9,117 | 8,770 | 7,940 |  |

**Supplementary Table 4:** Gene families unique only to P. downsi

|  |  |
| --- | --- |
| EGT47331 | hypothetical protein CAEBREN_08836 [Caenorhabditis brenneri] |
| GBO13109 | hypothetical protein AVEN_233885-1 [Araneus ventricosus] |
| GBP44057 | PiggyBac transposable element-derived protein 4 [Eumeta japonica] |
| KMQ83311 | transposable element tc3 transposase [Lasius niger] |
| KNC20639 | hypothetical protein FF38_06891, partial [Lucilia cuprina] |
| KNC20966 | hypothetical protein FF38_09018 [Lucilia cuprina] |
| KNC26906 | hypothetical protein FF38_10930 [Lucilia cuprina] |
| KXJ68891 | hypothetical protein RP20_CCG001210 [Aedes albopictus] |
| OAF68034 | hypothetical protein A3Q56_04227 [Intoshia linei] |
| PCG68510 | hypothetical protein B5V51_5157, partial [Heliothis virescens] |
| XP_011295145 | PREDICTED: tigger transposable element-derived protein 6-like isoform X2 [Musca domestica] |
| XP_011295374 | PREDICTED: tigger transposable element-derived protein 6 [Musca domestica] |
| XP_013109704 | PREDICTED: uncharacterized protein LOC106088638 [Stomoxys calcitrans] |
| XP_017475229 | PREDICTED: uncharacterized protein LOC108365650 [Rhagoletis zephyria] |
| XP_017478109 | PREDICTED: uncharacterized protein LOC108367917 [Rhagoletis zephyria] |
| XP_017478991 | PREDICTED: uncharacterized protein LOC108368617 [Rhagoletis zephyria] |
| XP_017479715 | PREDICTED: uncharacterized protein LOC108369194 [Rhagoletis zephyria] |
| XP_017481195 | PREDICTED: RNA-directed DNA polymerase from mobile element jockey-like [Rhagoletis zephyria] |
| XP_019891578 | PREDICTED: ATP-binding cassette sub-family A member 3-like [Musca domestica] |
| XP_019894716 | PREDICTED: uncharacterized protein LOC105262305 isoform X1 [Musca domestica] |
| XP_021704105 | protein ALP1-like [Aedes aegypti] |
| XP_022823959 | piggyBac transposable element-derived protein 4-like [Spodoptera litura] |
| XP_022834134 | uncharacterized protein LOC111361914 [Spodoptera litura] |
| XP_033325321 | uncharacterized protein LOC117219890 [Megalopta genalis] |
| XP_036214104 | trypsin zeta-like [Bactrocera oleae] |
| XP_036337749 | uncharacterized protein LOC118747737 isoform X3 [Rhagoletis pomonella] |
| XP_036342696 | uncharacterized protein LOC118751975 [Rhagoletis pomonella] |

**Supplementary Table 5:** Number of genes in *Fibrinogen C-Terminal Domain-Containing* gene family and *SCP domain-containing* gene family

| Species | Fibrinogen C-Terminal Domain-Containing gene family | SCP domain-containing gene family |
| --- | --- | --- |
| *H.melpomene* | 1 | 2 |
| *A.aegypti* | 33 | 20 |
| *C.capitata* | 10 | 6 |
| *S.calcitrans* | 45 | 17 |
| *M.domestica* | 32 | 25 |
| *P.downsi* | 5 | 7 |
| *G.morsitans* | 7 | 7 |
| *D.melanogaster* | 11 | 15 |

**Supplementary Figure 1:**
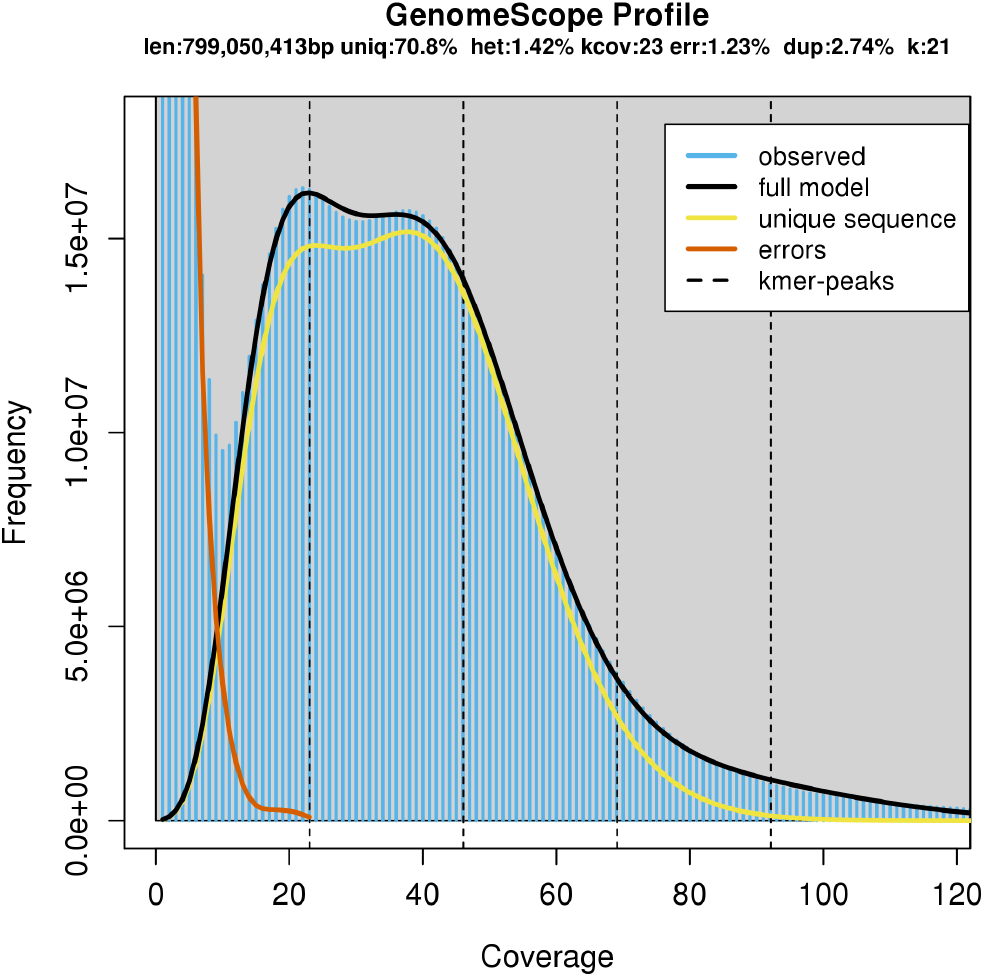
k-mer spectrum and fitted modelling used for estimating genome parameters of P. downsi from short sequencing reads

**Supplementary Figure 2:**
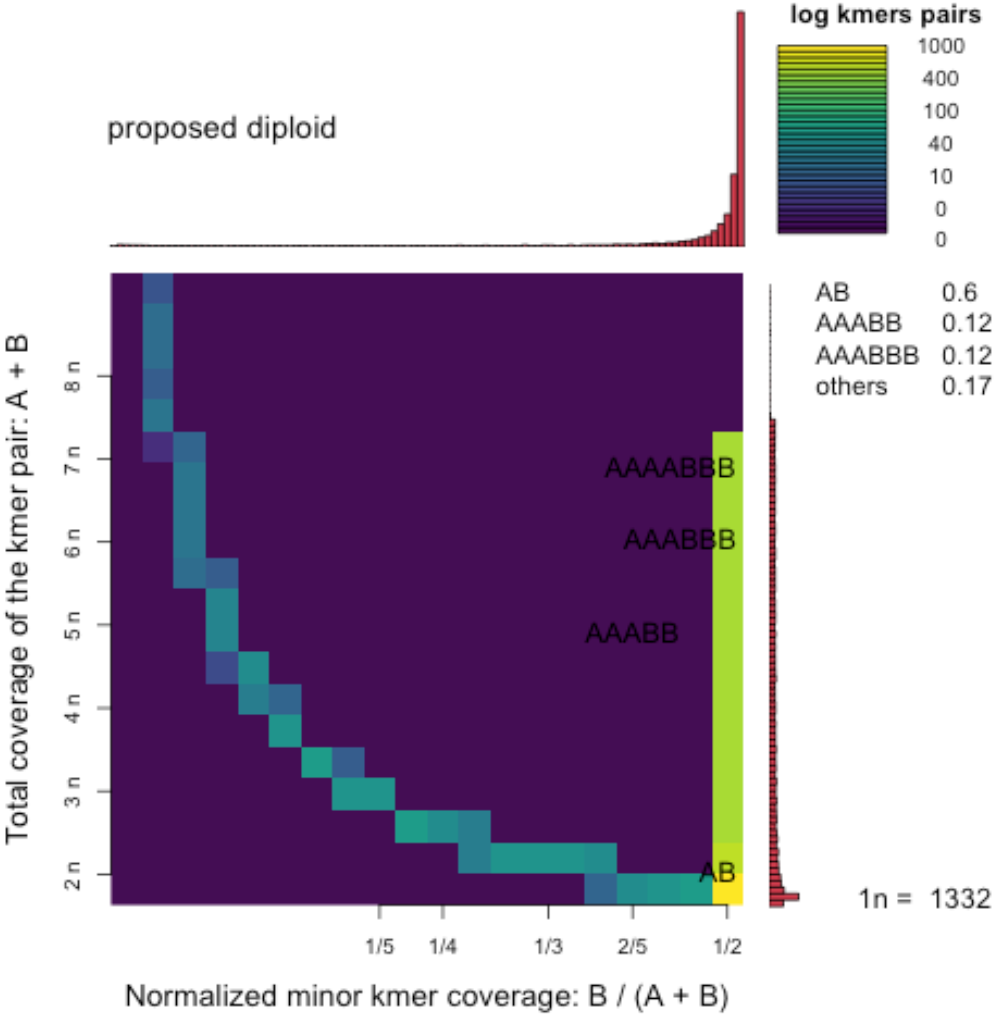
Distribution of the total coverage of the k-mer pair (y-axis) against relative minor k-mer coverage (x-axis) providing evidence of diploidy in P. downsi.

**Supplementary Figure 3:**
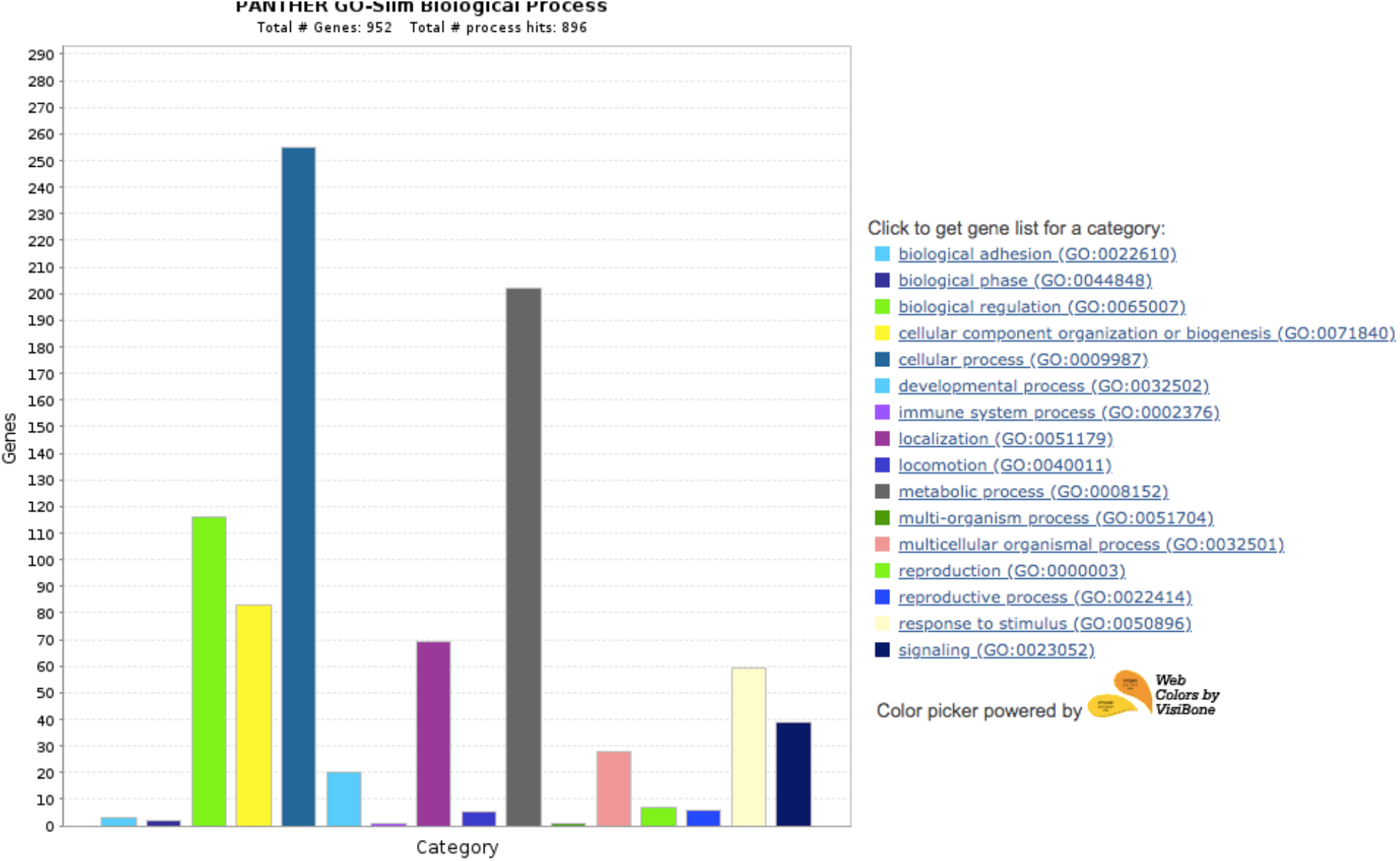
Gene ontology terms associated with 993 gene families unique to diptera

**Supplementary Figure 4:**
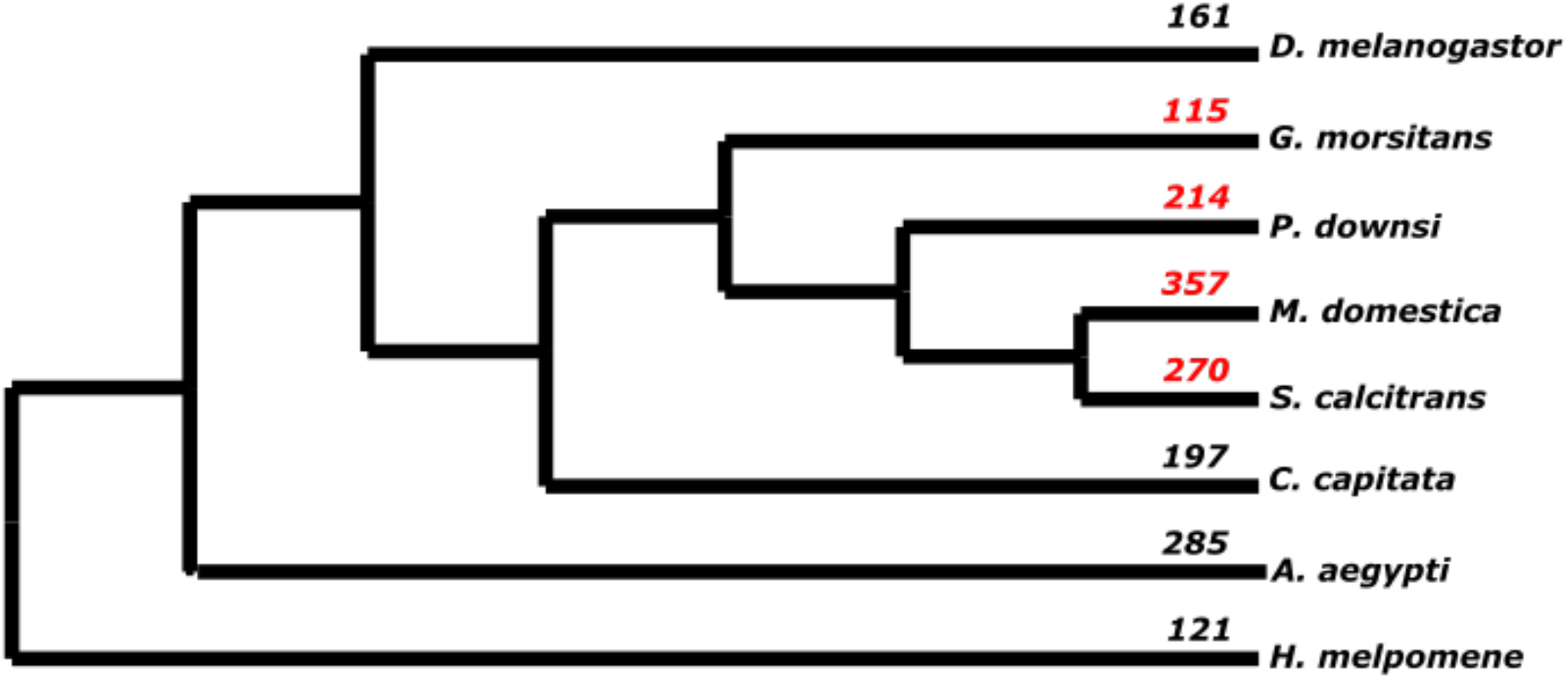
Number of genes in Cuticular gene family across dipterans

